# Hierarchical emergence of opponent coding in auditory belt cortex

**DOI:** 10.1101/2024.10.22.619743

**Authors:** Jeffrey S. Johnson, Mamiko Niwa, Kevin N. O’Connor, Brian J. Malone, Mitchell L. Sutter

## Abstract

We recorded from neurons in primary auditory cortex (A1) and middle-lateral belt area (ML) while rhesus macaques either discriminated amplitude-modulated noise (AM) from unmodulated noise or passively heard the same stimuli. We used several post-hoc pooling models to investigate the ability of auditory cortex to leverage population coding for AM detection. We find that pooled-response AM detection is better in the active condition than the passive condition, and better using rate-based coding than synchrony-based coding. Neurons can be segregated into two classes based on whether they increase (INC) or decrease (DEC) their firing rate in response to increasing modulation depth. In these samples, A1 had relatively fewer DEC neurons (26%) than ML (45%). When responses were pooled without segregating these classes, AM detection using rate-based coding was much better in A1 than in ML, but when pooling only INC neurons, AM detection in ML approached that found in A1. Pooling only DEC neurons resulted in impaired AM detection in both areas. To investigate the role of DEC neurons, we devised two pooling methods that opposed DEC and INC neurons – a direct subtractive method and a two-pool push-pull opponent method. Only the push-pull opponent method resulted in superior AM detection relative to indiscriminate pooling. In the active condition, the opponent method was superior to pooling only INC neurons during the late portion of the response in ML. These results suggest that the increasing prevalence of the DEC response type in ML can be leveraged by appropriate methods to improve AM detection.

## INTRODUCTION

Amplitude modulation (AM) is a fundamental feature of many ecologically relevant signals, including communication signals. The detection of AM is thus a fundamental task in the auditory system. Importantly for the understanding of neural coding, however, the representation of AM is profoundly transformed across the ascending auditory system. Arguably, the most salient aspect of this transformation is the reduction in synchronization to the modulation envelope along the ascending auditory neuraxis (1, 2), which limits the explicit encoding of AM in the firing patterns of auditory neurons. Nonetheless, it is possible that other aspects of this transformation could inform the encoding and decoding of AM signals.

The neural representation of AM is transformed from primary auditory cortex (A1) to other cortical areas (3–9) including area ML (10–13). From the standpoint of population coding, the two most important response changes are the reduction in synchronization and the increasing heterogeneity of rate tuning. Reduced response synchrony constrains the utility of spike timing-based modulation encoding at the level of individual neurons (14–19) and neural populations (20, 21), while the increase in the diversity of rate-based tuning constrains the effectiveness of pooling rate responses (22).

Most A1 neurons increase their firing rates at higher modulation depths (10). By contrast, a substantial portion of neurons in the middle lateral (ML) field of the belt region of auditory cortex decrease their firing rates when the depth of the modulation applied to a carrier increases (11). When a given brain region includes neurons that exhibit (predominantly monotonic) increases and decreases in responsiveness for increases in a given stimulus parameter (e.g., modulation depth), we refer to this representation as “bidirectional encoding”. Although decreasing responses to increasing modulation depths can be observed in A1, they are relatively uncommon (e.g., 11, 15). Decreasing responses approach parity with increasing responses in field ML, which suggests that this representation is an emergent property of the ascending auditory pathway. This change in the relationship between firing rate and modulation depth implies that the optimal readout mechanism for A1 and ML populations should differ. Here, we investigate how differences in A1 and ML cortical responses affect modulation depth detection by simulating a number of distinct population-based decoding methods that rely on spike rates, or response synchrony. Specifically, we test the hypothesis that a “two-pool opponent code” leverages the emergent diversity in rate tuning for modulation depth observed in ML (13).

Population codes are highly relevant to models of auditory processing because they allow for more complex sensory representations than single neuron-based encoding models, due to the exponential increase in the dimensionality of the population response space as the size of the population increases (23–26). Because neural populations can, and typically do exhibit high heterogeneity both within and across structures, they can encode stimulus information in a wide variety of ways (27–29) and would thus require a similarly wide variety of downstream readout mechanisms to support effective decoding. Our goal here is to investigate how the representational changes from primary (e.g. A1) to secondary (e.g. ML) auditory cortex impact the utility of broad classes of population-based decoding models.

In the current study, we explore a series of population-based decoding methods based on equal-weighted pooling of cortical neurons to quantify neurometric performance for modulation detection. Our results demonstrate that the increased prominence of bidirectional encoding in ML can be exploited by a two-pool opponent code, indicating that the optimal readout for higher-order auditory cortex is likely to differ from optimal readout methods for earlier auditory stages where the stimulus representation is distinct. These findings also suggest that the emergence of bidirectional encoding in the auditory belt might reflect an increasing need for stimulus representations that are robust to stimulus-extrinsic influences on firing rates related to task performance.

## MATERIALS AND METHODS

### Subjects

We made single neuron recordings from the right hemispheres of three adult rhesus macaque monkeys (V, female, age 11-12; W, female, age 10-14; X, male, age 13-16). We recorded from primary auditory cortex (A1) in all three animals (V, 22 cells; W, 144 cells; X, 126 cells). We recorded from the middle lateral belt area (ML) in two animals (W, 99 cells; X, 74 cells). Recordings analyzed in this study were made in conjunction with recordings analyzed for prior publications. Data from passive modulation transfer function (MTF) collection were not used in this study but have been reported previously (13), and some data from active and passive test blocks were included in previous publications (10–12, 30, 31). The analyses presented in this study have not previously been published. All procedures conformed to the United States Public Health Service (PHS) policy on experimental animal care and were approved by the UC Davis animal care and use committee.

### Identification of cortical fields

The identification of the cortical field location of each recording site was based on physiological measurements and has been described in detail for these recordings in Niwa et al. (10, 11). A tonotopic map collected with pure tones was used to identify A1. The A1/ML border was determined on the basis of differences in tone latency, frequency tuning width, and tone/bandpass noise preference. Area ML was 2-3 mm wide and located lateral to the A1/ML border. The anterior border of ML with the anterolateral belt area (AL) and the posterior border of ML with the caudolateral belt area (CL) were estimated using a tonotopic map of best frequency (BF) determined with a bandpass noise tuning curve.

### Stimulus generation and presentation

The acoustic stimuli were 800-ms sinusoidally amplitude-modulated (AM) non-bandlimited Gaussian noise bursts with a 100-kHz sampling frequency. The stimulus set included modulation frequencies (MFs) of 2.5, 5, 10, 15, 20, 30, 60, 120, 250, and 500 Hz, and modulation depths of 0% (unmodulated), 6%, 16%, 28%, 40%, 60%, 80%, and 100%. For all stimuli, the appropriate modulation envelope was applied to an identical “frozen” noise stimulus generated by a single random number sequence.

Stimuli were created in Matlab (MathWorks) with a 5-ms 1-cosine^2^ ramp at onset and offset. Digital stimuli were converted to an analog signal with a D/A converter (Power1401; Cambridge Electronic Design) at a sampling rate of 100 kHz, and then passed through both a programmable attenuator (PA5; Tucker-Davis Technologies) and a passive attenuator (LAT-45; Leader) before amplification (MPA-200; Radio Shack) and delivery to a speaker inside the sound booth. Two different sound booths (IAC) with different speakers were used. One booth was 2.9 × 3.2 × 2.0 m with a Radio Shack PA-110 speaker (10-inch woofer with piezo-horn tweeter) positioned 1.5 m in front of the animal; the other booth was 1.2 × 0.9 × 2.0 m with a Radio Shack Optimus Pro-7AV speaker positioned 0.8 m in front of the animal. In both booths the center of the speaker was positioned at ear level. Stimulus intensity was calibrated to 63 dB sound pressure level using a sound level meter (model 2231; Brüel *&* Kjær) while the animal was absent with the microphone placed in a position corresponding to the center of the animal’s head when present.

### Behavioral procedure

Animals were trained to discriminate noise-carrier sinusoidally modulated AM (AM noise) from unmodulated noise. The experiments reported here were comprised of three different experimental trial-block types: [1] a passive (no task) modulation transfer function (MTF) block; [2] active test blocks; and [3] passive (no task) test blocks. In an initial MTF block [1], we collected an MTF using 100% modulated stimuli. The MTF of the multiunit response was used to determine the multiunit best modulation frequency (BMF) for both rate (spike count, *SC*) and temporal (phase-projected vector strength, *VS_PP_*, see below) measures. Either the rate or the temporal BMF (both based on multiunit activity) was pseudo-randomly selected as the test MF for subsequent active [2] and passive [3] blocks at varying modulation depths. The behavioral condition (active or passive) for the first test block was selected pseudo-randomly. During active test blocks [2], the animal was required to initiate each trial by holding down a lever. After a 500-ms delay, two 800-ms sounds were presented with a 400-ms interstimulus interval. The first sound (S1) was always unmodulated noise. The second sound (S2) was either unmodulated noise (non-target trials; ∼12.5% of trials) or modulated noise at one of the seven modulation depths (target trials; ∼87.5% of trials). The animal was required to listen through the offset of the second stimulus, then release a lever (“hit”) within an 800-ms response window on target trials or continue holding the lever on non-target trials (“correct rejection”). During passive test blocks (3), no S1 sounds were presented, and the animal was neither required to initiate the trial, nor to make a response. Because cells could not always be held long enough to complete a recording session, the number of trials per modulation depth varied. Active test blocks averaged ∼47 trials per depth; passive test blocks averaged ∼51 trials per depth. Some active recordings included in the analysis (A1: *n* = 94; ML: *n* = 26) do not have a passive counterpart because the cell was lost. Passive recordings without an active counterpart were excluded from further analysis.

To encourage alertness and good task performance, the animals were fluid regulated during the period of the study. Animals were rewarded with juice or water for correct responses (hits and correct rejections) during the active blocks. Passive blocks had occasional randomized liquid presentations.

### Physiological recording

Each animal was implanted with a titanium head post and head-restrained during recordings in a custom-made “acoustically transparent” wire mesh primate chair. Access to A1 and ML was provided by a recording chamber (CILUX, Crist Instruments) located over parietal cortex, and containing a plastic grid with 27-gauge holes arranged in a 15 × 15 mm square (1 mm intervals). Before each recording, a stainless steel guide tube was inserted through the grid and transdurally into the brain. A high-impedance tungsten microelectrode (1–4 MΩ, FHC; 0.5–1 MΩ, Alpha-Omega) was inserted into the guide tube, and a hydraulic microdrive (FHC) was used to lower the electrode through parietal cortex into A1 or ML.

The electrophysiological signal was amplified (A-M Systems model 1800) and filtered (0.3-10kHz; A-M Systems model 1800 and Krohn-Hite model 3382) then sampled at 50 kHz by an A/D converter (Power 1401; Cambridge Electronic Design) and saved to hard disk. Spike sorting of action potentials was performed offline (Spike2; Cambridge Electronic Design). Spike sorting was implemented by a waveform matching algorithm that represented candidate spike waveforms in 3-D PCA space. Well-isolated clusters in PCA space were defined as single units. A k-means clustering algorithm (Spike2; Cambridge Electronic Design) was applied to overlapping clusters in PCA space; waveform templates were created from the k-means clusters when the centers matched the centers of density in PCA space.

### Data analysis

All data analysis was performed using Matlab (MathWorks). All statistical comparisons between counts or percentages were performed with a z-test for independent proportions. All other statistical comparisons were performed with a two-sided t-test.

### Measure of phase locking

To measure phase locking we used phase-projected vector strength (*VS_PP_*), a trial-based measure of synchrony which penalizes vector strength (*VS*) values on a given trial by their deviation from the cell’s overall mean phase (17). The standard formula for vector strength is

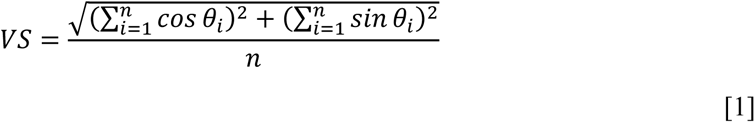

where *VS* is the vector strength, *n* is the number of spikes, and θ*_i_* is the phase of each spike in radians, calculated by

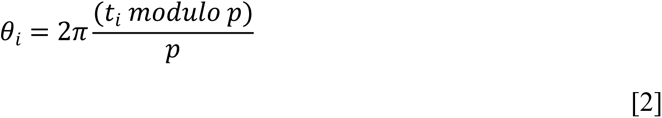

where *t_i_* is the time of the spike in ms relative to the onset of the stimulus and *p* is the modulation period of the stimulus in ms (32, 33).

Phase-projected vector strength (*VS_PP_*) was calculated on a trial-by-trial basis as follows:

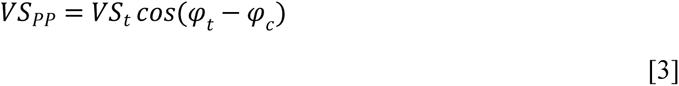

where *VS_PP_* is the phase-projected vector strength per trial, *VS_t_* is the vector strength per trial, calculated as in Eq. 1, and φ*_t_* and φ*_c_* are the trial-by-trial and mean phase angle in radians respectively, calculated for each stimulus condition as follows:

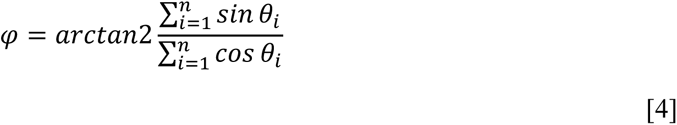

where *n* is the number of spikes per trial (for φ*_t_*) or across all trials (for φ*_c_*) and arctan2 is a modified version of the arctangent that determines the correct quadrant of the output based on the signs of the sine and cosine inputs (Matlab, ***atan2***). A cell that fired no spikes was assigned a *VS_PP_* of zero.

### Classifying modulation depth functions and calculating neurometric detection thresholds

We determined neurometric depth sensitivity for 259 cells in A1 and 143 cells in ML. Because some cells were tested at two modulation frequencies, we had a total of 292 recordings in A1, and 173 recordings in ML. Cells were tested at the multiunit-based rate BMF, the multiunit-based temporal BMF, or both, when the recording duration allowed. We employed modulation depths of 0% (i.e., unmodulated noise), 6%, 16%, 28%, 40%, 60%, 80%, and 100%. We calculated the area under the receiver operating curve (ROC area) (34) at each nonzero modulation depth. The ROC area measures how well the neural responses on each trial can be used to distinguish modulated and unmodulated stimuli. ROC area was computed for both *SC* and *VS_PP_*. ROC area integrates to 1 if all of the values in the modulated distribution are greater than all of the values in the unmodulated distribution, and 0 if all of the values in the unmodulated distribution are greater than all of the values in the modulated distribution. In both cases, an ideal observer would predict with 100% accuracy whether or not a given spike train was elicited by a modulated stimulus. An ROC area of 0.5 corresponds to chance performance.

We define ROC areas plotted across modulation depth as modulation depth functions (MDFs) for each recording: a rate-based MDF using spike count (rMDF) and a temporal-based MDF using phase-projected vector strength (tMDF) were created. A given cell can indicate the presence of modulation by increasing the value of some metric (for example firing rate) calculated on its responses or by decreasing that value. We determined whether each MDF belonged to an “increasing” (INC) or “decreasing” (DEC) response class by averaging ROC areas across modulation depth. MDF averages above 0.5 were classified as increasing; MDF averages below 0.5 were classified as decreasing. This means for an MDF classified as increasing, on average, the firing rate is higher than the firing rate in response to unmodulated noise. For an MDF classified as decreasing, on average, the firing rate is lower than the firing rate in response to unmodulated noise. MDFs calculated from responses within a window from 400-800 ms after stimulus onset were used to determine whether a cell belonged to the increasing or decreasing response class for the purposes of the pooling analysis (see Results). In all other analyses, the response from the entire stimulus duration (i.e., 800 ms) was used.

To determine a modulation depth detection threshold (“threshold”) for *SC* and *VS_PP_* for each recording, we fit each rMDF and tMDF with two functions: 1) a logistic function [eq. 5], and, because MDFs need not be monotonic, 2) a Gaussian function [eq. 6]:

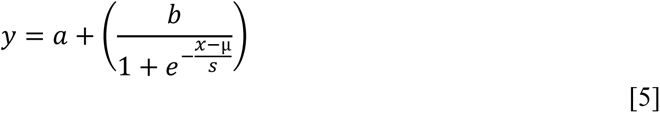

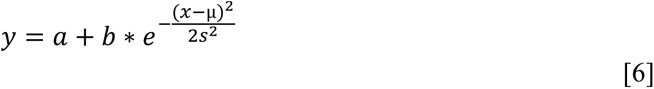

Both curves involve four free parameters: the y-offset (*a*), the height (*b*), the x-center (µ) and the slope (*s*). We used Matlab’s “fmincon” function to perform a constrained fit, where we restricted the slope (*s*) between 2 and 20 and restricted the height (*b*) to six times the difference between the ROC area value at 6% depth and the ROC area value at the most responsive (i.e., furthest from 0.5) modulation depth. To prevent overfitting, we did not fit Gaussian functions [eq. 6] if the absolute ROC area value (|ROC area – 0.5|) at 100% depth was more than 7/8 of the maximal absolute ROC area value of the MDF. If both fits were attempted, we used the fit with the highest correlation coefficient with the MDF. The threshold was taken as the modulation depth at which the fitted function crossed (or first crossed in the case of a Gaussian) the value of 0.75 for an increasing function, or the value of 0.25 for a decreasing function. If this criterion was not crossed, the cell was deemed not to have reached detection threshold.

### Pooling

For our pooling analyses, we pooled spikes from multiple cells on a trial-by-trial basis before performing subsequent analyses. These multi-cell spike trains were composed of the union of the spike times across individual cells as follows:

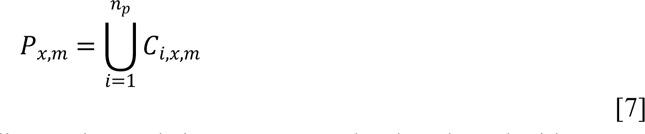

where a number of neurons corresponding to the pool size *n_p_* were randomly selected with replacement from all neurons (in all animals) tested at modulation frequency *m* (excluding 500 Hz due to low sample size) from a population *x*, which was defined by 1) cortical area (A1 or ML); 2) behavioral condition (active or passive); 3) response class (increasing, decreasing, or all); and 4) spike train epoch (all, “early” [0-400 ms], or “late” [400-800 ms]). Each randomly selected cell had its trial order at each modulation depth randomly scrambled to generate cell *C_i,x,m_*, which had 50 trials at each modulation depth. Randomization of trial order ensured that the probability that the same trial was pooled with itself in any *P_x,m_* was low when a particular recording was randomly selected multiple times in the same *P_x,m_*. For recordings with more than 50 trials at a given modulation depth, the first 50 scrambled trials were included. For recordings with fewer than 50 trials, the balance of trials was taken from a re-randomization of the original set of trials. Thus, each *P_x,m_* was a list of 50 pooled trials (built from *n_p_* cells) at each modulation depth. ROC area for *SC* and *VS_PP_* was calculated on the pooled trials in each *P_x,m_* and a pooled modulation detection threshold was calculated by fitting the ROC areas with equations [5] and [6] as above. This procedure was repeated 1000 times for each *n_p_*. The percentage of pools reaching threshold was defined as the “success rate”. The mean threshold of pools (that managed to reach threshold) was calculated for each *n_p_* and for each tested MF. For most analyses in this study, these values were averaged across MF for presentation. We employed pool sizes of *n_p_* = 1, 2, 3, 4, 5, 6, 7, 8, 10, 12, 16, 25, and 50 units.

We performed two variants on the above pooling technique to investigate whether the existence of increasing and decreasing MDFs could be leveraged to increase success rates and decrease pooled modulation detection thresholds. In “subtractive” (“SUB”) pooling, spikes from decreasing MDFs have an inhibitory (*ergo* subtractive) effect on the pool. For SUB pooling, separate *P_x,m_* were built for the increasing and decreasing MDFs at each *n_p_*. Since the selection of cells was random, it was possible for the increasing *P_x,m_* or decreasing *P_x,m_* to be empty, particularly for low pool numbers. For *SC* calculations, the spike count on each pooled trial was equal to the spike count from the increasing *P_x,m_* minus the spike count from the decreasing *P_x,m_*, with a floor of 0. The floor was used to model the property that neural firing rates don’t go below 0. For *VS_PP_* the SUB model was designed to be sensitive to whether spikes from increasing rMDFs were in phase or counterphase to those from decreasing rMDFs (there are no decreasing tMDFs as neurons do not phase lock to 0% modulation). For these calculations the pooling process directly eliminated spikes; there was no subtraction of firing rates. For each spike in the decreasing *P_x,m_*, we deleted the first spike (if present) within 5 ms of the “inhibitory” spike in the corresponding pooled trial in the increasing *P_x,m_*. *VS_PP_* was then calculated on the resulting pooled-and-eliminated spike trains. Calculation of ROC area, fitting of MDFs, and determination of threshold were then performed as described above.

The second variant employed a form of opponent coding, which we termed two-pool opponent coding (OPP), based on the notion that populations that increased and decreased firing rate in response to AM could be directly compared in a push/pull fashion (e.g., 35, 36). For OPP coding, an increasing *P_x,m_* and a decreasing *P_x,m_* were independently created at each *n_p_*, as in normal pooling, and *SC*/*VS_PP_* was calculated on each pooled trial in each *P_x,m_*. The calculated value for each trial in the decreasing *P_x,m_* was then subtracted from the calculated value for its corresponding trial in the increasing *P_x,m_* to create the OPP *P_x,m_*. ROC area, MDF fitting, and threshold determination was performed on the OPP *P_x,m_* as described above.

For *SC*, OPP pooling differed from SUB pooling in two ways: 1) The response classes were balanced, so INC and DEC spike counts had equal weight. By contrast for SUB pooling, we used random selection, which will give more weight to the more abundant response class. 2) OPP pooling did not have a floor of zero, to simulate a “push-pull” mechanism, so if a given pair of DEC and INC pools had more DEC spikes, the pooling metric was allowed to be negative. For *VS_PP_*, OPP pooling differed from SUB pooling because *VS_PP_* values were calculated independently for INC and DEC pools before subtraction. Pool size for OPP was defined to be the summed total of cells in the two pools, resulting in OPP pool sizes ranging from 2-100, rather than 1-50.

### Behavioral thresholds

To facilitate comparison to the physiological results, we calculated behavioral thresholds from the active conditions of the recordings used in this study. Because thresholds were similar across animals, and because our pooling analyses pooled recordings across animals, we combined behavioral results from all three animals before calculating behavioral thresholds. For each modulation frequency, we calculated the signal detection theoretical measure *d’* for each modulation depth. The resulting *d’* measures for each depth were fitted with a standard logistic function [Eq. 5, above], and the threshold depth was taken as the point on the fitted function where *d’* = 1.

## RESULTS

We recorded amplitude modulation depth functions in A1 (259 cells, 33 of which were tested at two MFs for a total of 292 recordings) and ML (143 cells, 30 of which were tested at two MFs for a total of 173 recordings).

There was a lot of diversity in A1 and ML neural responses to noise-AM. Figures 1 and 2 show examples for four neurons capturing a large range of responses. The upper panels contain raster plots where phase locking and response magnitude can be seen, and the lower panels show neurometric depth versus ROC plots and curve fits for the calculation of AM detection threshold based on ROC areas.

**Figure 1:**
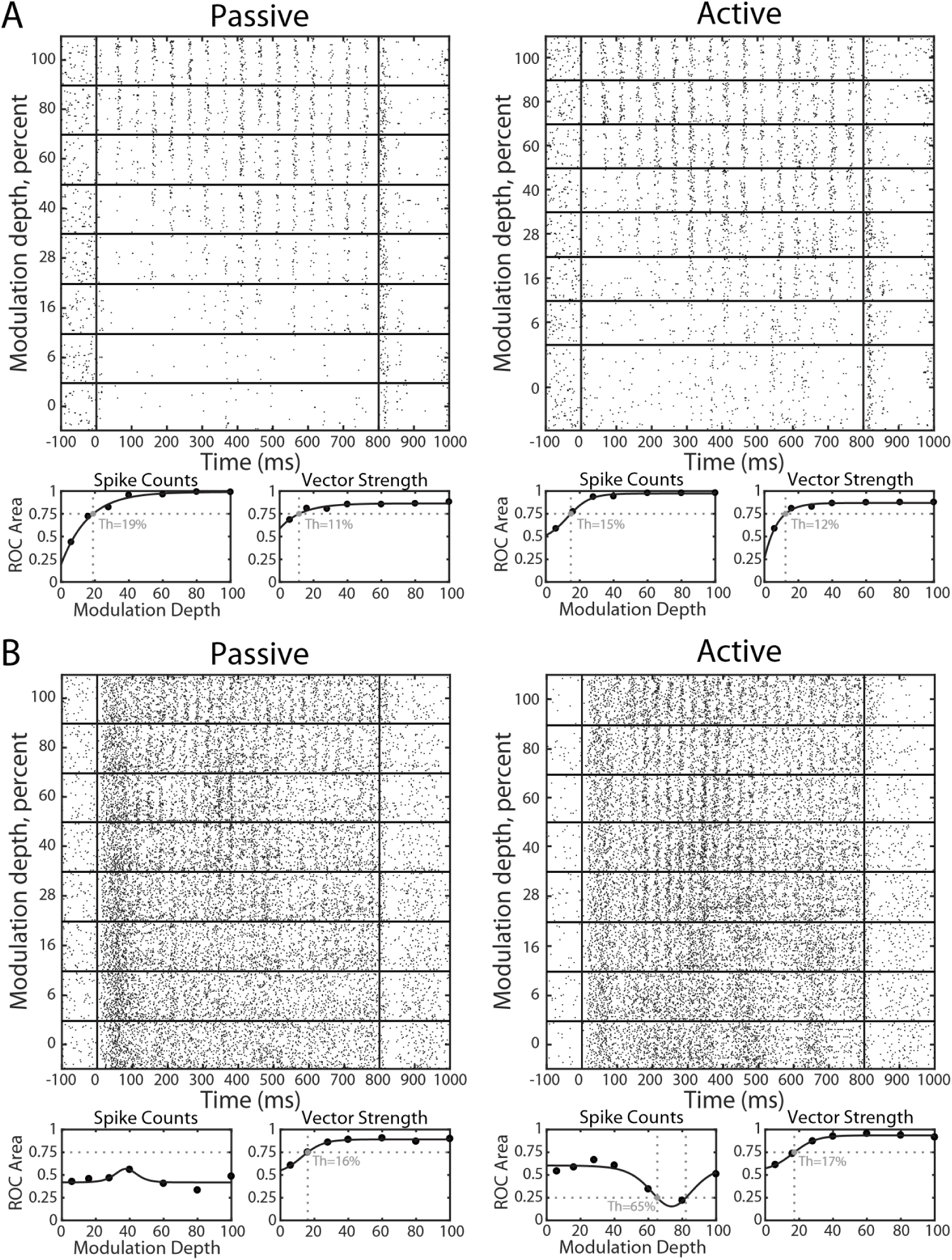
Reponses to AM in two example neurons. A: Spike train rasters and the associated MDFs depicting the responses of a neuron in A1 stimulated with 20 Hz AM applied to a noise carrier in the passive (left) and active (right) behavioral conditions. Vertical lines on raster plots indicate the beginning and end of stimulus presentation. Horizontal lines on raster plots separate the tested modulation depths. ROC Area is plotted against modulation depth (dots, with fitted line) for Spike Counts and phase-projected Vector Strength (*VSPP*). Vertical dashed gray lines indicate the modulation detection threshold (listed on each panel), the intersection point with the threshold value (horizontal dashed gray lines; 0.75 for increasing MDFs; 0.25 for decreasing MDFs). B: Responses of a neuron recorded in ML stimulated with 30 Hz modulation. Details as in (A).

**Figure 2:**
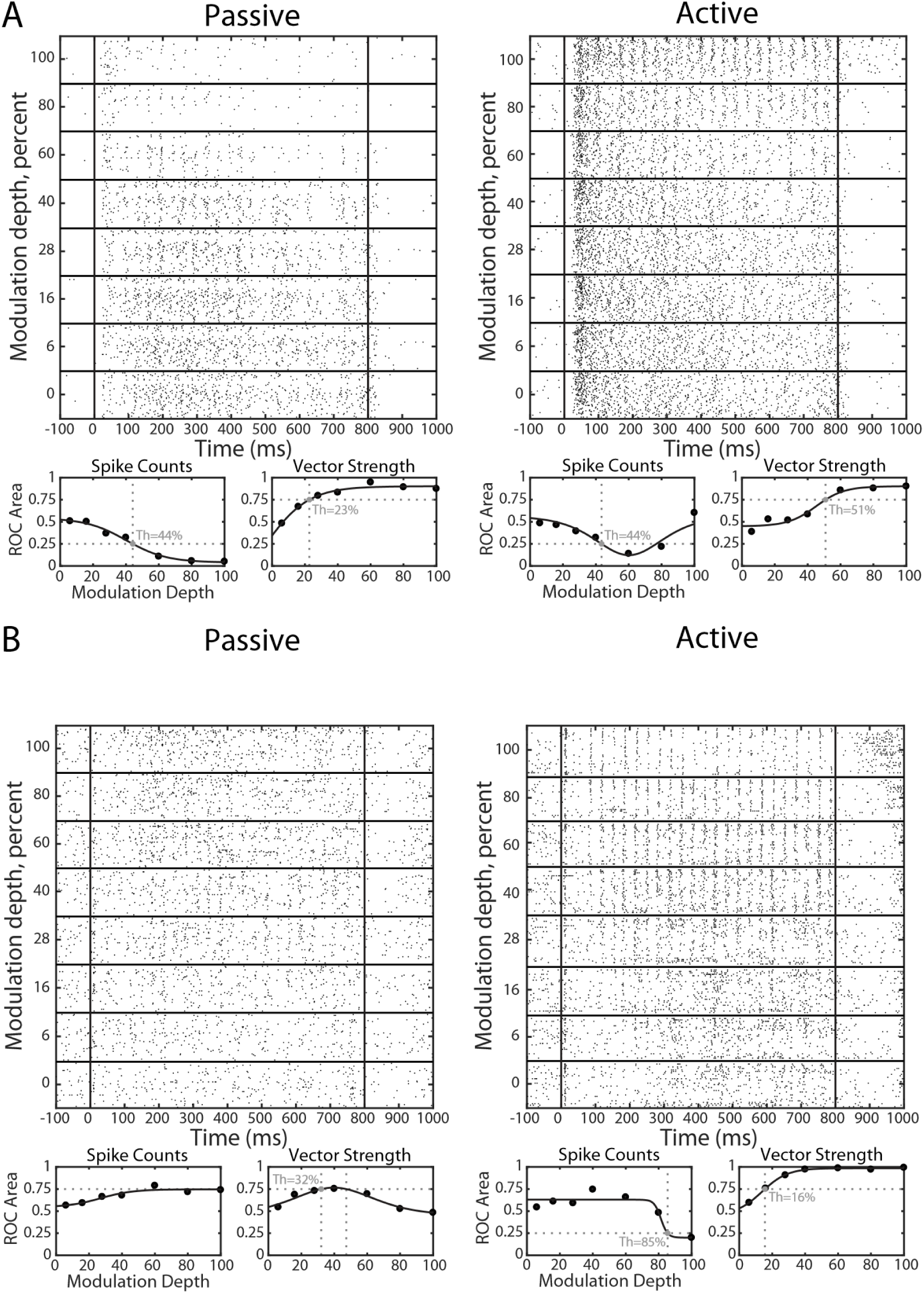
Reponses to AM in two atypical neurons. Plot details are the same as Fig. 1. A: Responses of a neuron recorded in A1 tested with 30 Hz modulation. B: Responses of a neuron recorded in ML, tested with 30 Hz modulation.

Figure 1A shows raster plots and MDFs of a neuron that phase locked well both in the passive and active conditions. Strong phase locking was common in A1. This cell fires more spikes during the active condition than the passive condition which also was quite typical (31). The neuron has a slightly lower *SC*-based neurometric AM detection threshold in the active than the passive condition (15% versus 19%), and very similar *VS_PP_*-based neuromeric thresholds across active and passive condition (12% and 11%). This was more sensitive than the typical A1 neuron, but its improvement in average *SC*-based ROC-area thresholds in the active condition and lack of significant difference between active and passive conditions for *VS_PP_*-based ROCa are typical of A1 cells (31).

Fig. 1B depicts responses from an ML neuron. In the passive condition, this cell has no defined *SC*-based threshold, but in the active case, the spike count decreases observed at 60 and 80 percent modulation depth cross the threshold (ROCa = 0.25) for distinguishing AM noise from unmodulated noise. Despite a higher proportion of non-phase locked spikes relative to the example in Fig. 1A, this neuron has worse but roughly comparable *VS_PP_*-based ROCa AM detection thresholds (16% and 17%).

The A1 neuron with responses depicted in Fig. 2A has strongly suppressed responses at the highest modulation depths in the passive condition, but not in the active condition. This was atypical. Despite the strong suppression of activity at higher modulation depths in the passive condition, the spikes that did fire at these higher depths were very temporally precise causing strong phase locking and a monotonically increasing *VS_PP_* vs MDF (lower right panel in Fig. 2A). In both active and passive conditions, the firing rate for modulated stimuli is typically lower than that for unmodulated stimuli, resulting in ROC areas less than 0.5 for this decreasing neuron.

Fig. 2B shows responses of a neuron in ML where the diversity of responses was even more apparent than in A1. This cell is unusual because its phase locked responses disappear at high modulation depths in the passive condition, but not in the active condition, where phase locking is remarkably tight (recall that the *VS_PP_* -based MDF depicts ROC area, not actual *VS_PP_* values). Typically, A1 phase locking was stronger and improved monotonically with modulation depth. Overall, Figs. 1 and 2 are consistent with the idea that neuron response characteristics were diverse rather than stereotypical.

### Mean modulation depth functions

As an initial characterization of our data, we averaged MDFs across modulation frequency for the active and passive conditions. We included only those cells that were tested at the same modulation frequency in both the active and passive conditions. For spike count (*SC*) mean MDFs, we separated our cells into increasing (INC) and decreasing (DEC) response classes. Because Niwa et al. (11) showed large changes in activity during the second half of stimuli (400-800 ms) in the active condition for some cells, we used only the second half of the response in the active condition to define the INC and DEC response classes in this study, rather than the full stimulus duration. In A1, this change resulted in a total of 24 cells (8% of the total) changing response class and a net shift of 10 cells into the DEC class (217 INC [74%], 75 DEC [26%]). In ML, this change resulted in a total of 34 cells (20% of the total) changing response class, and a net shift of 10 cells into the DEC class (95 INC [55%], 78 DEC [45%]) relative to classifications based on the full stimulus duration.

Mean MDFs, plotted by active/passive condition (green/orange) are shown in Figure 3. *SC*-based MDFs (Fig. 3A-D) are separated into INC and DEC response classes and by cortical area (A1: left; ML: right). *VS_PP_*-based MDFs for INC and DEC response classes are very similar, and so are only separated by cortical area (Fig. 3E-F). Active and passive mean MDFs were qualitatively similar. For *SC*, there is a trend for the active condition to have a higher firing rate than the passive condition, but this difference is significant only for 100% modulated stimuli in the INC response class in A1 (Fig. 3A), and for the spontaneous rates in A1. Mean firing rates for the INC response class are higher in A1 than in ML (three-factor ANOVA, *p* < 10e-8); mean firing rates for the DEC response class are also higher in A1 than in ML (three-factor ANOVA, *p* < 10e-15). Phase locking in A1 is stronger than in ML (three-factor ANOVA, *p* < 10e-6), consistent with the well-documented loss of phase locking in higher auditory centers (1, 2).

**Figure 3:**
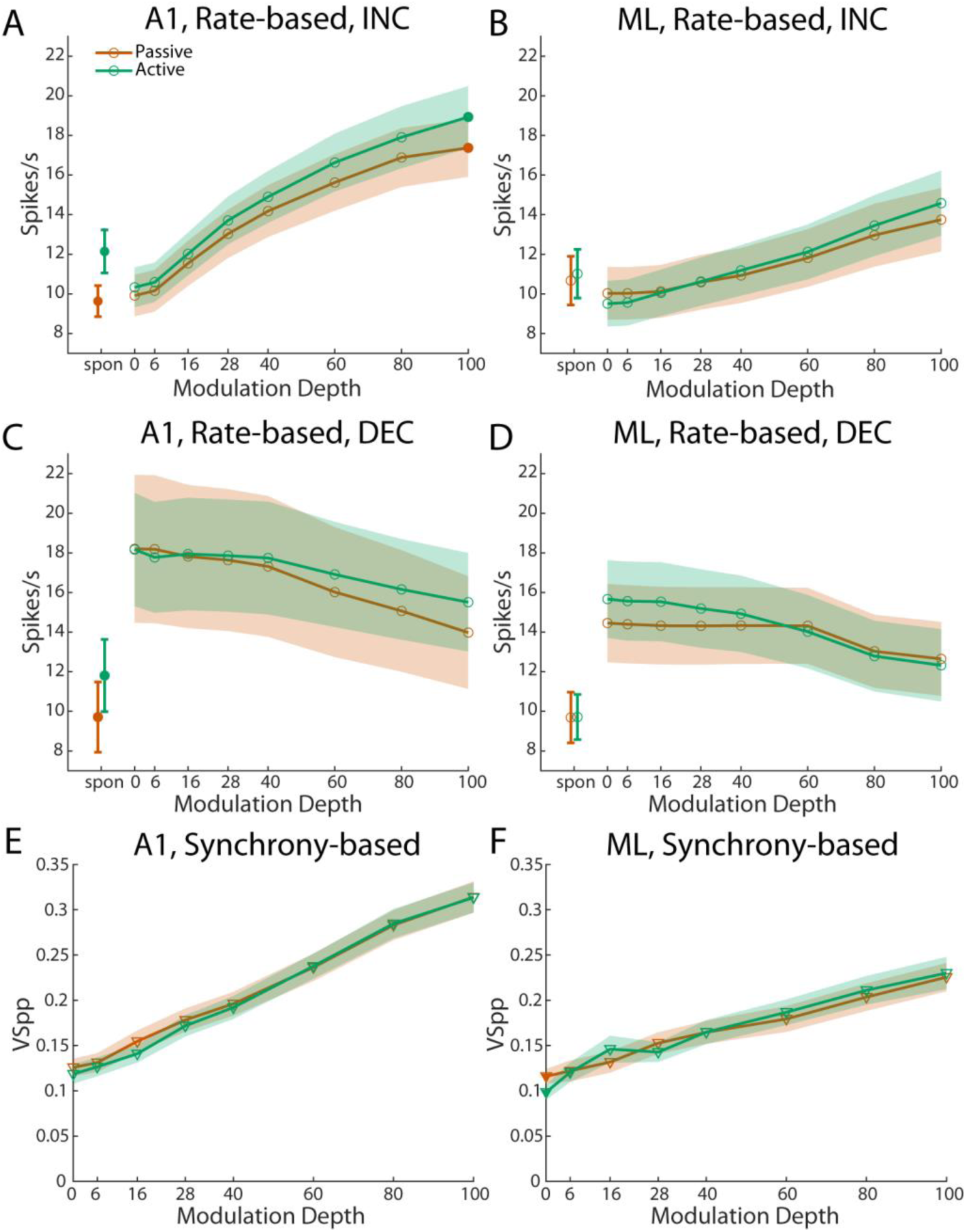
Mean modulation depth functions. Mean MDFs were calculated from the MDFs of cells tested at the same modulation frequency in both the active and passive conditions. Filled symbols indicate data points where active and passive conditions significantly differed (paired t-test). Error bars and shaded areas indicate +/- standard error of the mean. A, B: Mean spike rate MDFs for increasing cells in A1 (A; n = 142) and ML (B; n = 77). Spontaneous period is 100 ms prior to onset of stimulus. Spontaneous activity in A1 (p = 4.7e-5) and 100% depth in A1 (p = 0.045) differed significantly between the active and passive conditions. No other active/passive comparisons were significant. C, D: Mean spike rate MDFs for decreasing cells in A1 (C; n = 48) and ML (D; n = 65). Spontaneous period is 100 ms prior to onset of stimulus. Spontaneous activity in A1 (p = 0.017) differed significantly between the active and passive conditions. No other active/passive comparisons were significant. E, F: Mean *VSPP* MDFs for A1 (E; n = 190) and ML (F; n = 142). No active/passive comparisons were significantly different.

### Indiscriminate pooling of responses in A1 and ML

It is probable that behavior is driven by the activity of ensembles of neurons rather than by single neuron responses alone. We investigated the ability of ensemble responses to detect amplitude modulation by performing offline pooling of our responses across separate recording sessions (see Methods). For one method, pooling was performed separately at tested MFs and consisted of randomized trial selection and randomized cell inclusion, with replacement, for population sizes from 1–50 neurons. We call this form of pooling “indiscriminate” because responses are pooled together without respect to any cell’s response properties. After pooling the data across a given population, we applied the same neurometric analysis used on single neurons in the lower panels of Figs. 1-2 to determine neurometric AM detection thresholds for the pools.

Figure 4 illustrates the results for two different methods of evaluating the effect of response pooling on AM detection. The top row shows the success rate, i.e., the percentage of pools that successfully detect modulation as defined by reaching an ROC area threshold criterion of 0.75 for increasing depth-rate functions or 0.25 for decreasing rate-depth functions (Fig. 4A-B). A success was achieved if the fit to the depth-rate function crossed threshold (more details on fits are in Methods). Success rate is agnostic as to whether the neuronal response has a low or high threshold with regard to the minimum modulation depth detectable by the pool. Success rate is depicted for both spike count (*SC*, left) and for vector strength (*VS_PP_*, right). Success rate increases smoothly as the number of pooled cells increases in all tested conditions and was consistently higher for A1-derived pools (red lines) than ML-derived pools (blue lines). *SC-* based pools are more likely to reach threshold in the active condition (triangles) than in the passive condition (squares).

**Figure 4:**
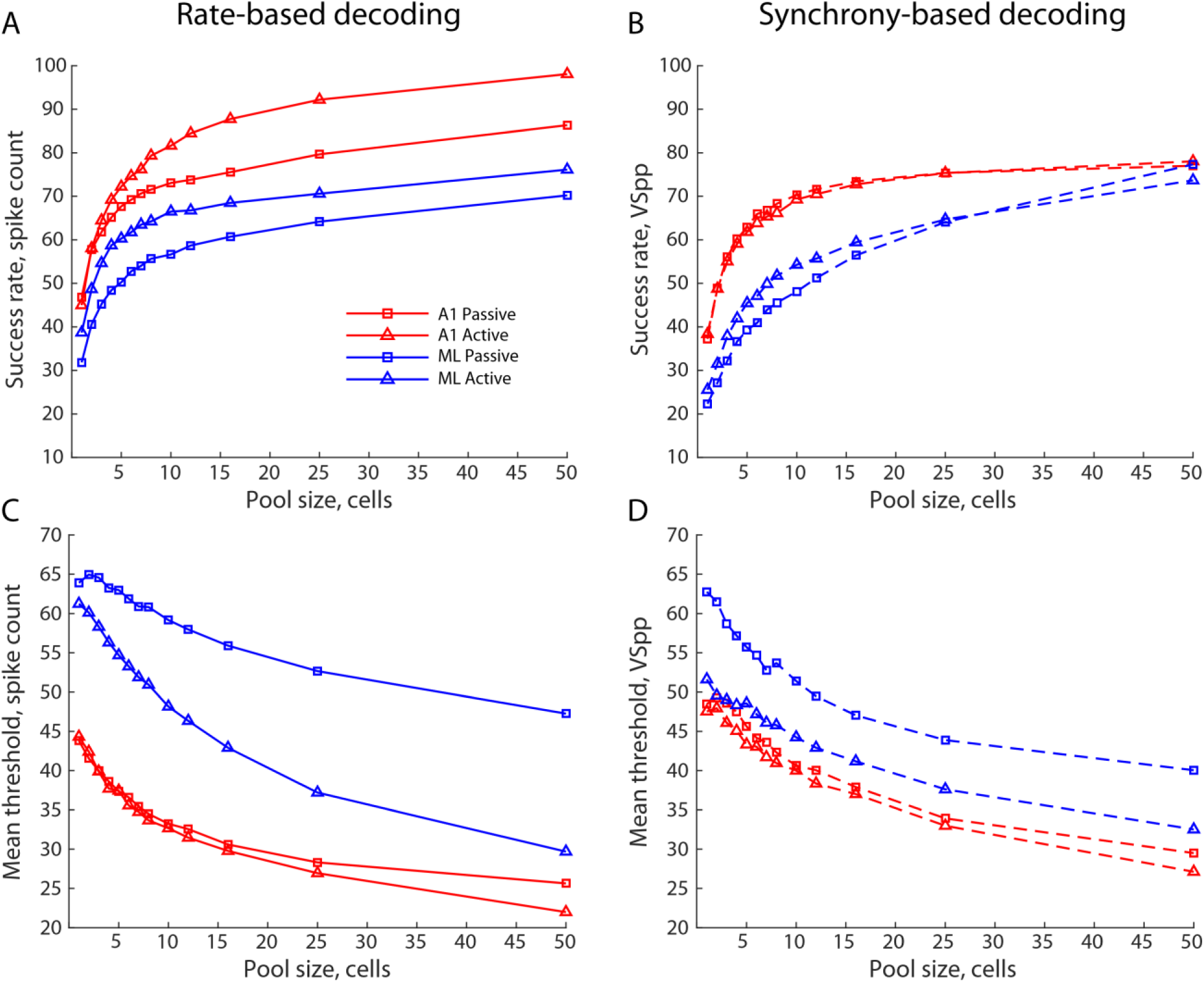
Indiscriminate pooling. A, B: Success rate for AM detection (defined in Methods as percentage of pools that reach threshold) with spike count (A) or vector strength (B) measures plotted against the pool size. Data are averaged across all tested modulation frequencies (MFs). C, D: Mean modulation depth threshold of pools that reached threshold for AM detection with spike count (C) or vector strength (D) measures plotted against the pool size. Data are averaged across all tested MFs (weighted by the number of pools reaching threshold at each MF).

We also evaluated the effect of pooling on AM sensitivity by calculating the mean modulation detection threshold *for all pools that reach threshold* (Fig. 4C-D). Mean thresholds decrease smoothly as the number of pooled cells increases for both *SC* (left) and *VS_PP_* (right), although occasionally there is a small increase in mean threshold for very small pools before the mean threshold begins to improve. As with success rate, A1-derived pools (red lines) have higher sensitivity (i.e., lower mean thresholds) than ML-derived pools (blue lines), and pools in the active condition (triangles) are generally more sensitive (lower mean thresholds) than corresponding pools in the passive condition (squares), particularly in ML-derived pools.

These data are replotted in a different format in Figure 5 to show statistical comparisons between different types of pools. The upper panels (Figs. 5A-B) depict A1-derived values plotted against their corresponding ML-derived values. In this and subsequent figures that depict data at different pool sizes where pool size is not a primary axis, symbol size indicates the pool size, with larger symbols corresponding to larger pools. The middle panels (Figs. 5C-D) depict active values plotted against their corresponding passive values; the lower panels (Figs. 5E-F) depict *SC*-derived values plotted against corresponding *VS_PP_*-derived values. Filled symbols indicate statistically significant differences (two-sided t-test, P < 0.05, corrected for multiple comparisons (i.e., 13 pool sizes) with respect to parity (dashed line) for the relevant axes on each panel set.

**Figure 5:**
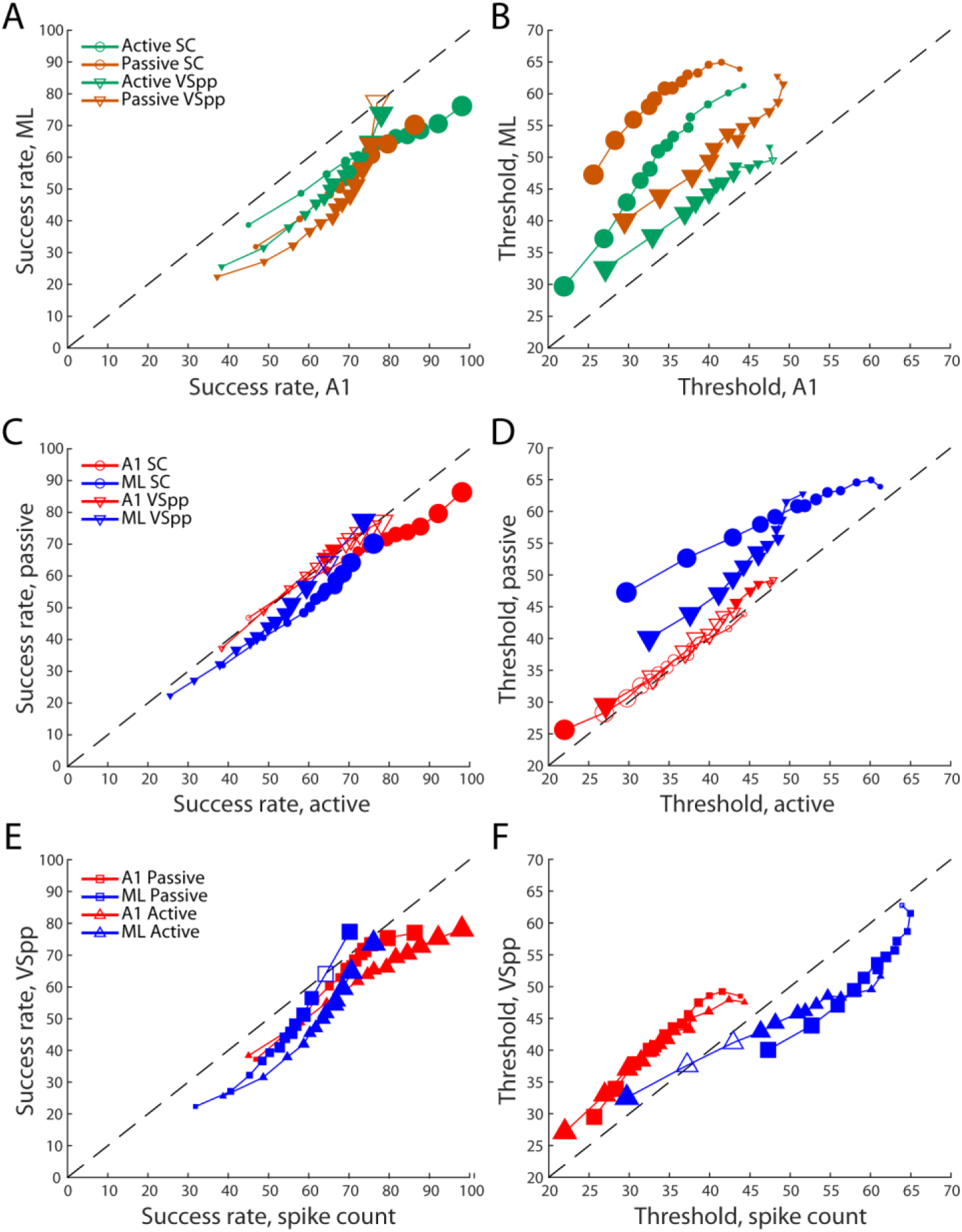
Indiscriminate pooling scatter plot. A: Comparison of success rates and thresholds between A1 and ML pools of neurons. This includes pools of all sizes from Fig. 4 with the symbol size in the scatter plot indicating the pool size, with larger symbols corresponding to larger pool A1 pools (x-axis) plotted against corresponding ML pools (y-axis). Symbol size increases with increasing pool size. Filled symbols indicate comparisons that are significantly different (two-sided t-test, P<0.05, corrected for 13 comparisons). Dashed line is a unity line. Pools below the unity line have a higher percentage of pools reaching threshold in A1 than in ML. B: As in (A), except plotting mean threshold of pools reaching threshold. Pools above the unity line are more sensitive (lower thresholds) in A1 than in ML. C, E: As in (A), except plotting active pools (x axis) against passive pools (y axis) in (C), and plotting spike count pools (x axis) against vector strength pools (y axis) in (E). D, F: As in (C, E), except plotting mean threshold of pools reaching threshold.

In all but one case, A1-derived pools outperform corresponding ML-derived pools, exhibiting higher success rates and lower mean thresholds for pools that reached threshold (since there is no threshold value for those that don’t). Active pools are generally more successful and more sensitive than corresponding passive pools, although there are three instances where the passive pool reached threshold more frequently than the corresponding active pool (where solid symbols are above the unity line on Fig. 5C), and there are quite a few non-significant comparisons for both success rate and mean threshold when comparing active and passive in A1 (panels C-D, red). The *SC*/*VS_PP_* comparison (Fig. 5E-F) shows a different pattern. *SC* pools exhibited higher success rates than corresponding *VS_PP_* pools in both A1 and ML (with one ML counterexample). However, *SC* measures resulted in lower mean thresholds than *VS_PP_* measures only for A1-derived pools. *SC*-derived thresholds were generally higher than *VS_PP_*-derived thresholds in ML-derived pools. This may seem surprising, given the reduction in *VS_PP_* in ML relative to A1, but these results must be considered in light of the fact that A1-derived pools exhibit lower thresholds than ML-derived pools for either response measure (Fig. 5B). More importantly, perhaps, A1 was dominated by the INC response class (74%), relative to the DEC response class (26%), whereas the ML categories were more closely balanced (55% and 45%, respectively), raising the possibility that pooling across response classes limited neurometric AM detection performance.

### Segregated pooling of increasing and decreasing response classes

In addition to simple indiscriminate pooling, we performed segregated pooling of INC and DEC response classes to test whether pooling these responses classes differently might affect AM detection. Figure 6 plots pools comprised of cells drawn from INC and DEC response classes against corresponding indiscriminate pools. Data points in Fig. 6 are collapsed across active and passive conditions. Filled symbols indicate exclusive INC (Fig. 6A-B) and DEC (Fig. 6C-D) pools that differ significantly from corresponding indiscriminate pools (two-sided t-test, P < 0.05, corrected for 13 comparisons).

**Figure 6:**
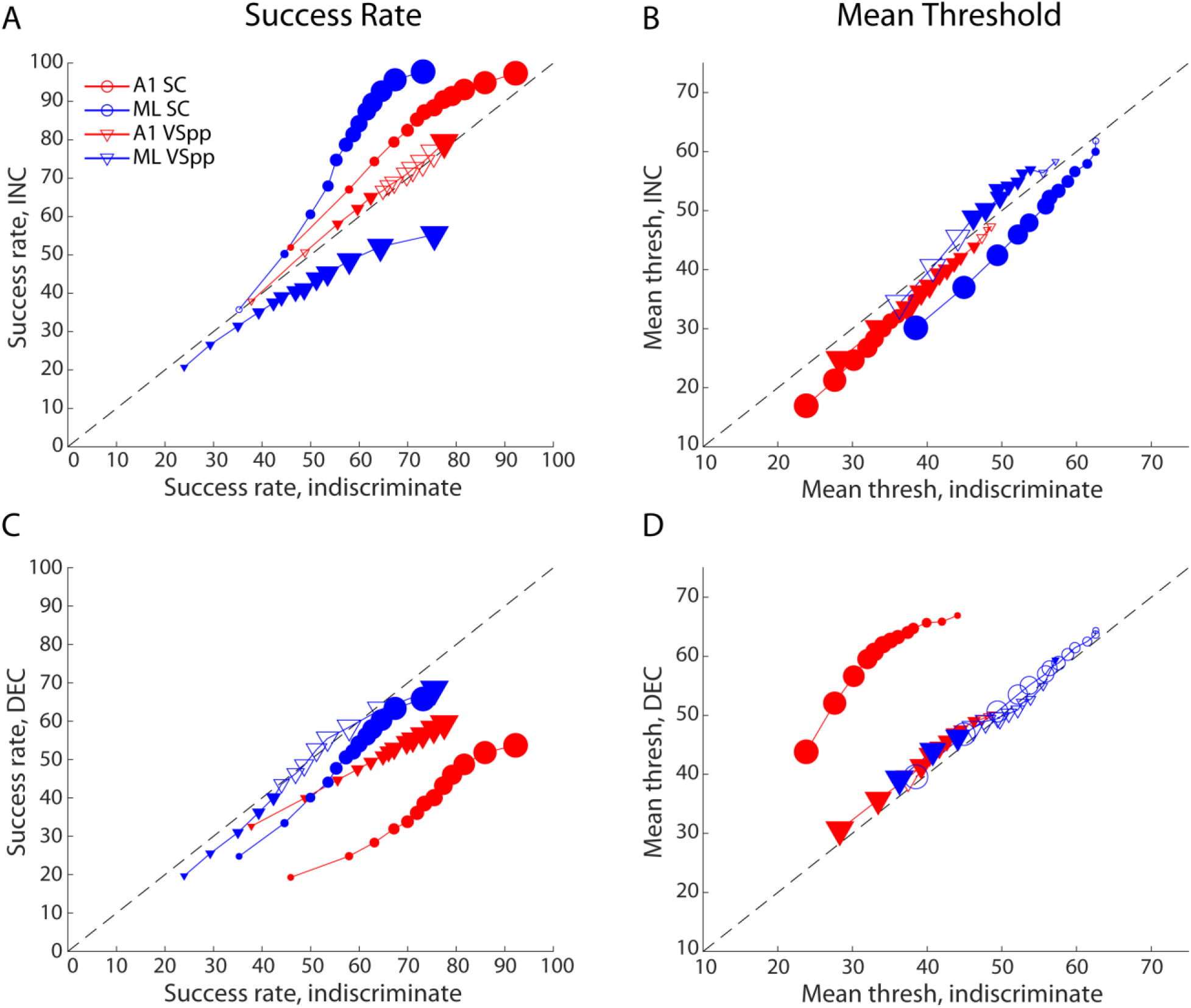
Segregated pooling of increasing and decreasing cells. All data points are averaged across active and passive conditions. For all plots, filled symbols indicate points where increasing/decreasing pools are statistically different than indiscriminate pools (two-sample t-test, P < 0.05, corrected for 13 comparisons). Symbol size increases with increasing pool size. A: Percentage of pools of increasing cells that reached threshold (y-axis) plotted against percentage of indiscriminate pools that reached threshold (x-axis). B: Mean AM detection threshold for pools of increasing cells (y-axis) plotted against mean AM detection threshold of indiscriminate pools (x-axis). C: As (A), except for decreasing cells. D: As (B), except for decreasing cells. INC, increasing; DEC, decreasing.

Comparing panels A and C, it is evident that success rates for INC pools generally exceeded those for DEC pools. The difference was particularly salient for *SC*-derived pools based on A1 responses (red circles), and ML responses (blue circles). This makes sense, since the definitions of the INC and DEC capture opposite changes in firing rate in response to AM. As a result, combining INC and DEC responses indiscriminately in a pool would diminish the apparent effect of AM. For A1, excluding roughly the quarter of DEC responses from the pools improves performance relative to indiscriminate pooling; for ML, excluding DEC responses (comprising 45% of the data sample) causes an even more pronounced improvement in success rate.

Results for the success rates of *VS_PP_*-derived pools exhibit a different pattern. There were slight improvements in the INC-only pool relative to indiscriminate for A1, but significant worsening for ML. It is important to recall that the response class definition (INC or DEC) was based on firing rate, not on response synchrony, so a class designation of DEC does not imply that *VS_PP_* values were reduced by modulation (in fact, they should not be, since for unmodulated controls *VS_PP_* should be near 0 as there is no modulation). As shown in Fig. 3E-F, *VS_PP_* values increased monotonically with modulation depth in both A1 and ML (INC and DEC responses were not segregated since they were roughly equivalent). For ML, removal of DEC responses from the pools lowered the *VS_PP_*-based success rate, as indicated by the values below the unity line (Fig. 6A; blue triangles).

The fact that the success rates were almost universally lower for DEC-only pools relative to indiscriminate pools suggests that DEC responses are typically less informative than INC responses (Fig. 6C). The predominance of INC responses in A1 is consistent with the large reduction in the *SC*-based success rate for A1 (red circles) and the smaller but significant reduction in *VS_PP_* -derived success rates (red triangles; compare Fig. 6A and 6C). The largest increase in the success rate occurs when limiting the ML-derived pools to INC responses (Fig. 6A, blue circles), as indicated by the distance from the unity line. Since INC and DEC responses were nearly balanced for ML, indiscriminate pools reliant on firing rate are likely (in the context of random selection) to be compromised, since superimposing AM on the carrier elicits both increases and decreases among the member neurons of the pool.

The most striking effect of limiting pools to pure INC or DEC responses occurred for *SC*-based mean thresholds in A1-derived pools (red circles; Fig. 6D), which suggests that AM-imposed reductions in firing rate are not reliably distinct from responses to the unmodulated carrier on a trial-by-trial basis. In ML, however, pools drawn entirely from DEC responses performed much like indiscriminate pools. By contrast, *SC*-based mean thresholds were lower for pure INC pools than for indiscriminate pools for both A1 and ML (Fig. 6B).

It is possible to gauge the relative informativeness of INC and DEC responses for A1 and ML by considering the relative plotted positions of pools of equivalent size for the different cortical regions. For example, success rates for *SC*-based, INC-only pools are quite similar; the relative leftward shift of the ML pools shows that ML performance is penalized to a greater extent by indiscriminate pooling. For DEC-only, *SC*-based pools, however, A1-derived pools have lower success rates and higher mean thresholds for all pool sizes, indicating that the DEC responses in ML support AM detection more reliably and more sensitively than those in A1. Nevertheless, DEC-only pools underperformed indiscriminately pooled cells (Fig. 6C), suggesting that INC responses capture AM more effectively overall, though the margin narrows from A1 to ML.

### Subtractive pooling and two-pool Opponent Coding

It is possible that information from INC and DEC response classes could be leveraged more effectively for AM detection than the additive pooling scheme described above. We implemented two additional methods: 1) Subtractive (“SUB”) pooling, and 2) two-pool opponent coding (“OPP”). In SUB pooling, pools were composed of cells randomly selected from the population without regard to response class, but DEC cells had an inhibitory rather than excitatory influence on the pooled response. For *SC*, the spike count from DEC responses was subtracted from the spike count from INC responses, with a floor of zero. For *VS_PP_*, spikes from INC responses and DEC responses were pooled separately, then for each spike in the DEC pool, the first spike in the INC pool within a 5 ms window was removed to simulate inhibition. If no spikes in the INC pool occurred within the 5-ms window, the DEC response spike had no effect.

The results of the SUB pooling are summarized in Fig. 7A-B. As in Fig. 6, data from the active and passive behavioral conditions are combined, unless the legend indicates otherwise. Success rates (Fig. 7A) were lower for SUB pooling than for indiscriminate pooling based on *SC* and *VS_PP_* measures for A1 and ML. Results based on mean detection thresholds were analogous (Fig. 7B): SUB pooling increased mean thresholds relative to indiscriminate pools, except for a few small pool sizes for *SC* in A1 and ML, and for *VS_PP_* in A1. Overall, these results indicate that the SUB population decoding model does not effectively leverage cortical firing patterns in either A1 or ML. If DEC responses provide inhibitory drive to a pooled AM detection mechanism, the readout mechanism is likely more sophisticated than the SUB model implemented here.

**Figure 7:**
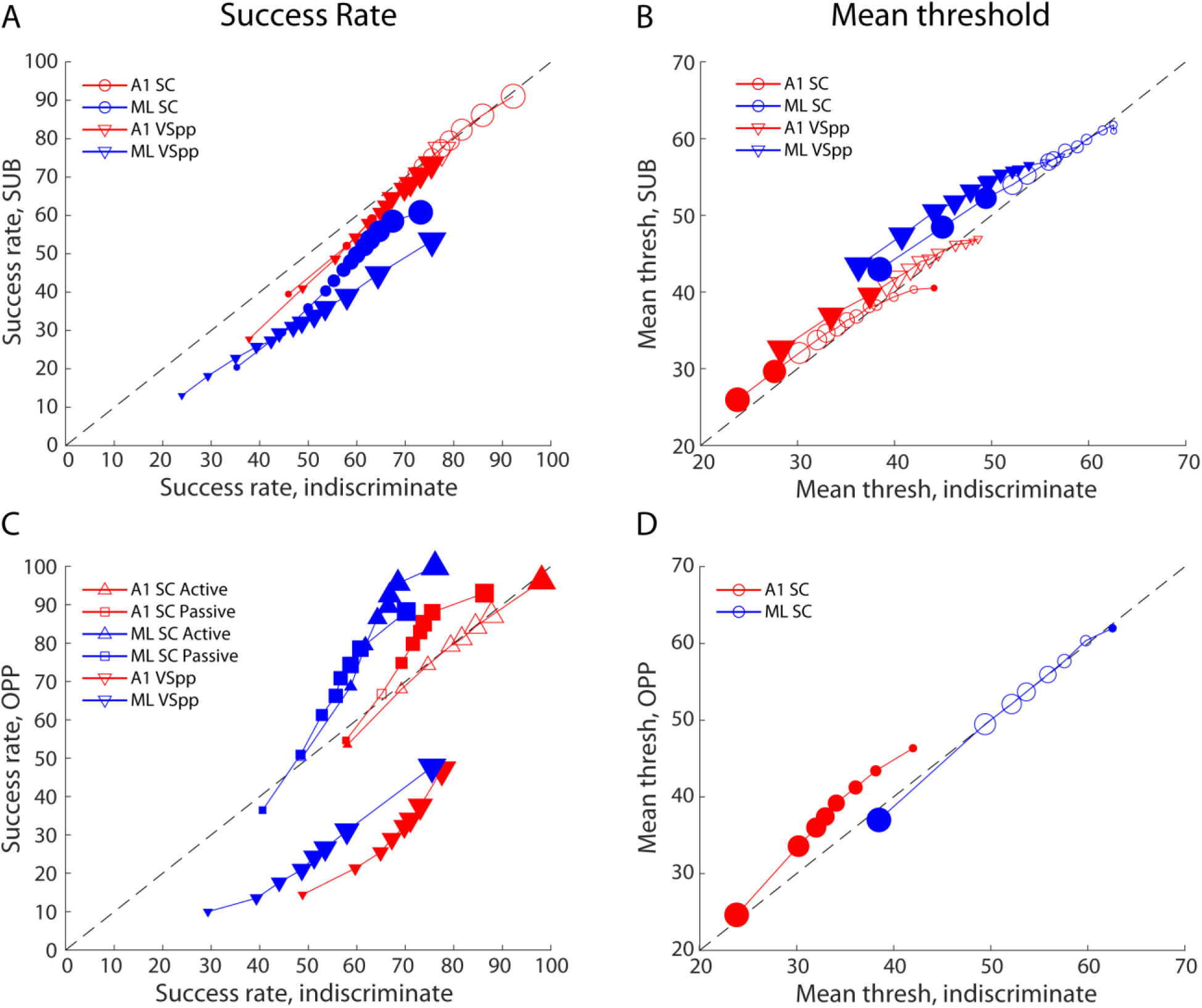
Combining increasing and decreasing cells. All data points are averaged across active and passive conditions unless otherwise noted. For all plots, filled symbols indicate points where increasing/decreasing pools are statistically different than indiscriminate pools (two-sample t-test, P < 0.05, corrected for 13 comparisons). Symbol size increases with increasing pool size. A: Percentage of pools using SUB pooling that reached threshold (y-axis) plotted against percentage of indiscriminate pools that reached threshold (x-axis). B: Mean AM detection threshold for SUB pools (y-axis) plotted against mean AM detection threshold of indiscriminate pools (x-axis). C: As (A), except for OPP; for *SC* measures, active and passive conditions are plotted separately. D: As (B), except for OPP pools. SUB, subtractive pooling; OPP, two-pool opponent.

In OPP pooling, two pools of equal size were generated from randomly selected INC and DEC responses, and *SC* and *VS_PP_* were calculated separately for each pool. The DEC pool result was subtracted from the INC pool result to obtain the metric used for neurometric analysis. For *SC*, OPP pooling differed from SUB pooling in two ways: 1) The responses classes were balanced, so INC and DEC spike counts had equal weight. By contrast, random selection for SUB pooling resulted in an average INC-to-DEC ratio of roughly 3-to-1 for A1 and 5-to-4 for ML; 2) OPP pooling did not have a floor of zero, to simulate a “push-pull” mechanism, so if a given pair of DEC and INC pools had more DEC spikes, the pooling metric was allowed to be negative. For *VS_PP_*, OPP pooling differed from SUB pooling because *VS_PP_* values were calculated independently for INC and DEC pools before subtraction. Pool size for OPP was defined to be the summed total of cells in the two pools, resulting in OPP pool sizes ranging from 2-100, rather than 1-50.

The results of OPP pooling are shown in Fig. 7C-D. Success rate (Fig. 7C) is depicted separately for active and passive conditions for *SC* measures because of notable differences between the two, particularly in A1. In the active behavioral condition, the OPP model applied to spike counts (*SC*) from A1 resulted in success rates very similar to those obtained with indiscriminate pooling (red “upward” triangles). In the passive condition, however, the OPP model outperformed indiscriminate pooling, particularly for larger pool sizes (red squares). When decoding spike counts from ML, the OPP model significantly outperformed indiscriminate pooling in both the active (blue “upward” triangles) and passive conditions (blue squares). The highest success rate we observed in this study occurred for the OPP model applied to ML data in the active condition using spike count as the metric. For a pool size of 100, 99.99% (8999/9000) pools reached threshold for AM detection.

As we expected, success rates for the OPP model based on *VS_PP_* (Fig. 7C; “downward” triangles) were much lower than those for indiscriminate pooling. We include the results here (combined across active/passive behavioral conditions) for completeness. Response classification (INC or DEC) was based on spike counts, because of the observation that AM can increase or decrease the firing rates of cortical neurons; there is no analogous effect for *VS_PP_*, which, on average, increases monotonically with modulation depth in both A1 and ML (Fig. 3E-F). In light of this, we did not further investigate the OPP models with *VS_PP_*.

Mean detection thresholds for OPP models applied to *SC* (averaged across active and passive conditions; Fig. 7D) did not exhibit the relative improvements we observed for success rates. In A1, OPP models resulted in higher mean thresholds than indiscriminate pooling; in ML, mean OPP thresholds were basically the same as for indiscriminate pooling, despite the increase in success rates. This discrepancy is likely explained by a recruitment effect: pools that do not reach threshold were not included in the average to obtain the mean threshold. The higher success rates for the OPP model added more high-threshold pools, thus raising the mean threshold.

### Separate pooling of early and late responses

Cortical responses to AM stimuli can evolve over time relative to stimulus onset, such that responses to the latter half of an 800 ms stimulus can differ from those in the initial half (11). For that reason, we analyzed “early” pooled responses (0-400 ms after stimulus onset) and “late” pooled responses (400-800 ms) separately. Figure 8 depicts the ratio (late/early) of the success rate (averaged across active and passive presentations) based on A1 and ML data for multiple decoding models: indiscriminate pooling, INC-only, and OPP (*VS_PP_*-based OPP models were excluded for reasons described in the previous section; see Fig. 7D). Success rates based on *SC* (Fig. 8A-B) generally suggest that late responses are more sensitive than early responses, except for indiscriminate pooling in ML, where early responses are somewhat more sensitive in all but the smallest pool sizes. *SC*-based late epoch increases in success rate were greatest for the OPP models in both A1 and ML. In ML, late epoch increases in success rate were evident for INC-only pools for all but the smallest pool sizes, whereas indiscriminate pools achieved higher success rates in the early epoch for all but the smallest pool sizes. In A1, by contrast, results were comparable for both models.

**Figure 8:**
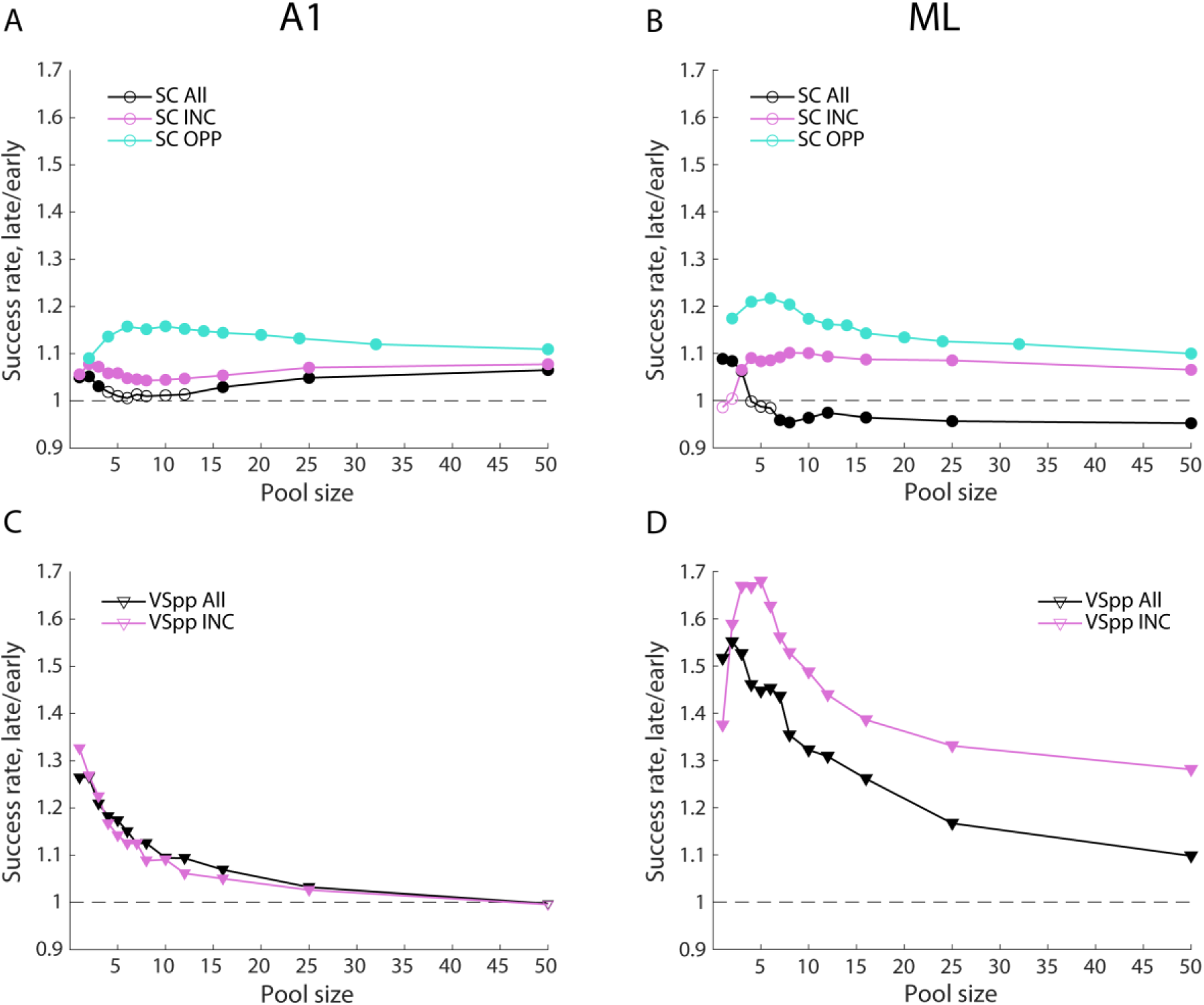
Comparing early and late responses. All data points are averaged across active and passive conditions. For all plots, filled symbols indicate points where “early” response (0-400 ms) pools are statistically different than “late” response (400-800 ms) pools (two-sample t-test, P < 0.05, corrected for 13 comparisons). A: Late/early ratio of number of pools reaching threshold (y-axis) plotted against pool size (x-axis) in A1 using *SC*. B: As (A), except in ML. C: As (A), except using *VSPP*. D: As (C), except in ML. All, indiscriminate pooling; INC, increasing; OPP, two-pool opponent.

The synchrony-based analysis (Fig. 8C-D) suggests that phase locking for individual neurons was enhanced in the late portions of the response, but that with larger pool sizes this effect dropped off. Most notably, success rates based on *VS_PP_* for ML show large relative increases during the late epoch for smaller pool sizes (Fig. 8D, downward triangles both magenta and black). Late epoch increases in success rate for *VS_PP_* were much smaller in A1, and limited to the smallest pool sizes (Fig. 8C, downward triangles both magenta and black).

We further refined our early/late epoch analysis to compare INC and opponent models because Figures 6 and 7 suggest that the largest gains in AM detection over simple indiscriminate pooling occurred for INC and for OPP pooling. This was particularly evident with success rate using *SC* measures. To investigate the relative performance of the OPP and INC models in greater detail, we computed ratios of the success rates subdivided by epoch and behavioral condition (Fig. 9). In A1 (red curves), INC-only pooling slightly outperformed the OPP model in the early epoch for both behaving and passive conditions (Fig. 9A). In the late epoch this was maintained for pools with less than 5 neurons. The larger pools in late epochs showed much smaller OPP/INC differences that, while significant, trended towards equality with increasing pool size.

**Figure 9:**
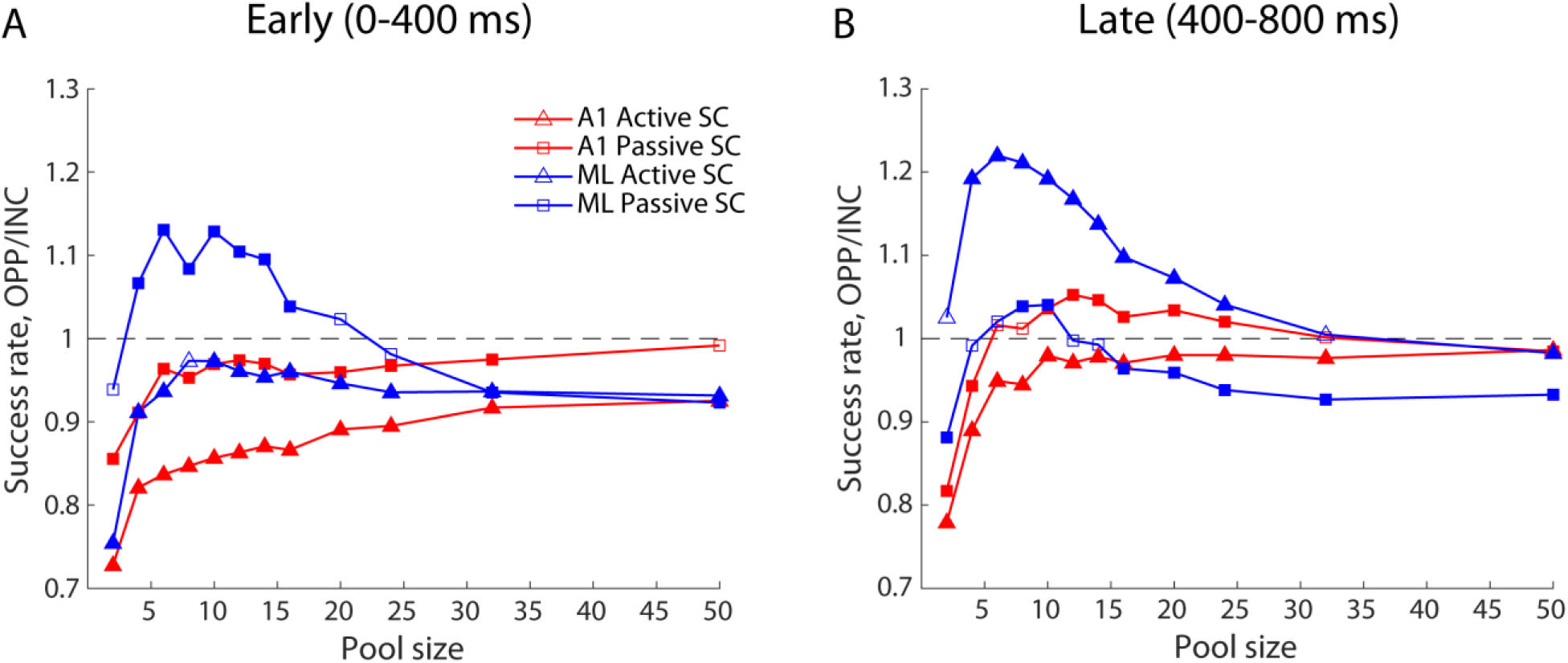
Comparing pools of increasing cells and two-pool opponent coding. For all plots, filled symbols indicate points where OPP pools are statistically different than INC pools (two-sample t-test, P < 0.05, corrected for 13 comparisons). A: OPP/INC ratio of number of pools reaching threshold (y-axis) plotted against pool size (x-axis) for early responses (0-400 ms). B: As (A), except for late responses (400-800 ms). INC, increasing; OPP, two-pool opponent.

In ML, however, there was a striking interaction between epoch and behavioral condition. In the late epoch (Fig. 9B), for the active condition (blue triangles), small OPP pools (*n* = 6-24) detected AM 20% more often than INC pools. By contrast, INC-only pools detected AM more reliably than the OPP model for larger pool sizes in the passive condition (blue squares). The opposite pattern was observed for the early epoch (Fig. 9A), where the OPP model outperformed INC-only pools in the passive condition (blue squares), and underperformed INC-only pools in the active condition (blue triangles). These results could explain the emergence of a large population of DEC responses in ML, since these responses could be leveraged, via a push/pull opponent code, to detect AM more reliably than INC responses alone.

### Comparison of pooling and behavioral results

Since population activity in auditory cortex likely drives the perceptual decisions underlying AM detection, we compared neurometric thresholds for the active behavioral condition against behavioral performance obtained during active recordings. The behavioral performance reported here closely matches that obtained for macaque monkeys engaged in the same psychophysical task, but collected with equal distribution across MF (37). The comparisons are plotted in Figure 10. Mean detection thresholds based on averages from single cells (squares) were substantially higher than behavioral thresholds (dashed lines) for all tested modulation frequencies for neurons in A1 and ML, regardless of the response metric (*SC*, *VS_PP_*).

**Figure 10:**
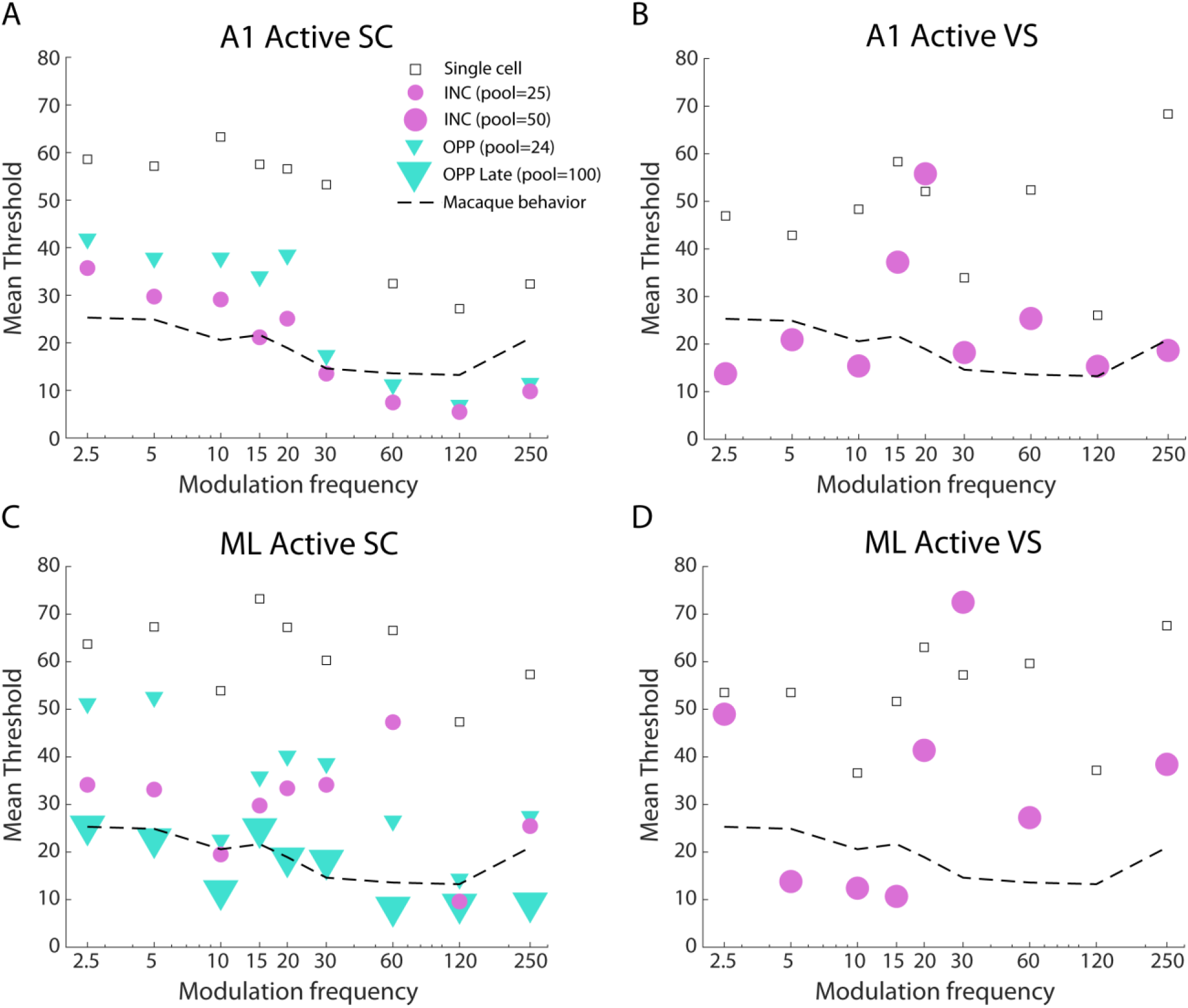
Comparing neurometric and behavioral thresholds. Behavioral thresholds for 800-ms AM stimuli are replotted from O’Connor et al. 2011. All data are from the active condition only. A: Behavioral and selected neurometric thresholds calculated with spike count (y-axis) plotted against modulation frequency (x-axis) for A1. B: Behavioral and selected neurometric thresholds calculated with vector strength (y-axis) plotted against modulation frequency (x-axis) for A1. C: As (A), but for area ML. D: As (B), but for area ML. INC, increasing; OPP, two-pool opponent.

Pooling spike counts from A1 resulted in significant improvements in AM detection thresholds. For example, INC-only pools based on 25 cells (small pink circles) resulted in neurometric thresholds much closer to the behavioral thresholds, although they underperformed at low modulation frequencies, and overperformed at higher modulation frequencies (Fig. 10A). In A1 relative to INC-only pools, the OPP model based on 24 cells (12 each in the increasing and decreasing opponent pools; small teal triangles) had higher thresholds at all MFs, particularly at modulation frequencies below 30 Hz. When relying on the *VSpp* metric (Fig. 10B), we found that a larger pool size (n = 50) provided the best fit to the behavioral performance, despite the poor fits at modulation frequencies of 15 and 20 Hz (compare with similarly poor *VSpp* performance in ref. 18, Fig. 9). In contrast to the results for *SC*, neurometric performance exceeded behavioral performance at the lowest modulation frequencies.

Results for ML-derived spike counts differed from those for A1 in a number of ways. First, mean thresholds for single cells were typically higher than those in A1 (compare Fig. 10A, C). Second, INC-only pools comprised of 25 cells did not approximate behavioral performance as closely as those in A1 (Fig. 10C; pink circles). Third, neurometric thresholds for the OPP model based on 24 cells (small teal triangles) typically underperformed the analogous neurometric thresholds in A1 (10 Hz was a notable exception). Lowest thresholds were obtained for a *SC*-based OPP model restricted to the late epoch (400-800 ms) of the response and including 100 cells (large teal triangles). This model also produced the best match to behavioral thresholds.

Similar to the results obtained in A1, *VS_PP_*-based decoding models for ML failed to capture behavioral thresholds effectively. INC-only pools resulted in neurometric thresholds far above the behavioral thresholds for all modulation frequencies other than a narrow range from 5 to 15 Hz, even for the largest tested pool sizes (n = 50). Given the overall patterns of neurometric thresholds in both A1 and ML, it would appear that *SC*-based pooling of cortical responses could plausibly account for the animals’ behavioral sensitivity, but *VS_PP_*-based pooling could not, unless the biological readout mechanism differs significantly from the decoding models we explored.

## DISCUSSION

Niwa et al. (10) first reported the existence of a significant minority of cells in ML that decreased their firing rates in response to increasing modulation depths, and based on that finding subsequently proposed that “separately decoding increasing and decreasing response functions as two complementary populations of neurons” might leverage that emergent response property in service of AM discrimination (11). Our current results indicate that an appropriate decoding model, a two-pool opponent code (OPP), can exploit decreasing modulation depth functions in service of modulation detection based on firing rates, particularly during task engagement. As Downer et al. (12) discussed, population-based neural codes allow for more complex sensory representations due to the increased dimensionality of the population response space for larger populations. In this paper, however, we explored lower dimensional subspaces of the response space by simulating convergence via response pooling, and by limiting the subpopulations to broad response-based categories (i.e., INC and DEC). The OPP code is effectively a two-dimensional decoding model, and its success in ML indicates that there exists a latent manifold of relatively low dimension that is adequate to account for behavioral performance, provided the pool size is sufficient (Figure 10); in A1, pooling limited to the dominant response type (INC) is sufficient for smaller pool sizes, given the higher sensitivity of individual A1 neurons.

The optimal readout for A1 and ML neurons differs because of the differences in the distributions of response properties in each field. The greater heterogeneity of ML responses disadvantaged rate-based pooling methods that did not segregate neuronal responses by tuning to modulation depth. This disadvantage was proportional to the relative parity of INC and DEC responses. In A1, DEC responses are far less common than in ML. As a result, the optimal pooling strategy for A1 was the exclusion of DEC responses, which contaminated the response pools, whereas for ML the optimal pooling strategy was an opponent code that exploited the greater prevalence of DEC responses. It is noteworthy that with the optimal readout strategy we explored for ML populations, the success rate for the two-channel opponent code was competitive with the optimal readout from A1, despite the reduction in synchrony and the increase in response homogeneity in ML (Figure 7). However, these results depend on both the analysis epoch (e.g. ‘early’ versus ‘late’) and the behavioral condition (‘passive versus ‘active’). Our results suggest that the optimal readout for a given population does not necessarily generalize across task demands, nor an animal’s training history.

The existence and prevalence of DEC cells indicates that the sign of the slope of cortical response functions can change dramatically from representations earlier in the auditory pathway, where monotonically increasing modulation depth functions are the norm. Interestingly, we did not observe representational changes that would indicate tuning for specific modulation depths, such as an increased prevalence of nonmonotonic tuning for specific, intermediate modulation depths. This is explicable in terms of the task demands, which required the animals to detect modulation at any depth, rather than discriminate among different depths.

The clear implication of our results is that any downstream readout mechanism must be at least as flexible as the underlying neural representation in order to support successful behavior. Moreover, prior work indicates that cortical responses are labile and adjust to task demands. In previously reported studies from our lab (12, 20, 38), animals were trained on a more complex task that required feature attention - the animals had to attend to either a change in spectral bandwidth or modulation depth. Under these task requirements, the distribution of DEC responses in A1 was quite different than that observed here - the proportion of A1 neurons that exhibited decreasing AM depth functions was more similar to that observed in ML in the current study. Moreover, the majority of A1 and ML neurons that exhibited significant AM depth encoding also exhibited increasing AM depth functions. In effect, the increases in attentional demands, stimulus feature ambiguity in firing rates, and task difficulty in the feature attention paradigm may have collapsed the distinction between A1 and ML observed in the current study. Based on these representational changes, the authors concluded that cortical AM encoding depends on behavioral and sensory demands.

Because in these studies an increase in either bandwidth or modulation depth could increase the firing rate of a given neuron, there is a potential confound for downstream neurons tasked with interpreting the perceptual basis for this change. Bidirectional encoding could aid in resolving this confound if there is a population of neurons that exhibit distinct changes in their response rates for increases in the behaviorally relevant stimulus features. This might explain the increased prevalence of DEC responses in A1 when the stimulus processing demands are more complex, as in the case of feature-based attention. If cortical feature representations are sufficiently malleable, we might expect, for example, that animals detecting a target at an intermediate modulation depth might develop nonmonotonic MDFs, rather than the largely monotonic MDFs (with positive or negative slopes) we observed in this and prior work (10, 13). In such a case, we expect that population-based readout mechanisms would show the requisite flexibility to exploit representational changes expressed in single neuron responses.

There are several important issues to consider with regard to the optimal readout of population responses in the context of highly trained animals performing a perceptual task. First, the optimality of the readout may depend on the attentive or behavioral state, and this effect may vary by cortical field. For example, the MDFs of neurons with decreasing responses in A1 showed shallower slopes during engagement relative to passive listening blocks; by contrast, neurons in ML with decreasing responses showed more steeply negative MDF slopes in the attending condition (11). Moreover, it cannot be assumed that the effects of attention are static and apply equally throughout stimulus presentation during each trial. For example, the differences in the relative performance of the OPP and INC decoding models in the early and late intervals relative to trial onset (Fig. 9) may be reflective of attentional shifts related to evidence accumulation and decision making (39, 40). These effects interact with pool size and vary by cortical field (Fig. 8), illustrating the complexity of the problem even in the context of population decoding models chosen for their simplicity.

With respect to an efficient or possibly optimal readout mechanism, we must emphasize that here we have employed fairly stringent limits on the population decoding models that we investigated in this study. Specifically, we only employed decoding models that simulated convergence via equal weighting of all neurons in a pool since we were interested in the potential implications of the representational changes we observed, particularly the bidirectional encoding of modulation depth. Individual weighting of neurons within each pool would have permitted the removal of divergent responses in that pool, such as the effective silencing of all DEC responses in A1 pools by setting their weights to zero. As Downer et al. (20) showed in the context of AM frequency discrimination, even a very small subset of the most individually informative neurons can achieve decoding performance approaching the performance for much larger pseudopopulations. Thus, improving the decoding model via individual weighting risks nullifying the existence of response categories by treating each individual neuron as its own effective type, as in the context of conventional labeled line codes (41–43).

The opponent code that proved most effective for rate-based population decoding in ML is distinct from—and arguably more sophisticated than—the subtractive model in two chief respects. First, the subtractive model enforces a floor of zero spikes, even when the simulated inhibition from pooled neurons with decreasing responses dominates excitation from neurons with increasing responses to larger modulation depths. Second, the opponent model balances the counts of increasing and decreasing responses in the opponent channels, while the subtractive model allows for imbalance among the pools since it is based on random draws. Thus, the opponent model implements a form of normalization absent from the subtractive model, and relaxes a rectification constraint, which benefits ML success rates (Fig. 7).

Notably, we did not observe similar patterns of improvement for the opponent model with respect to the mean thresholds for the pools. Success rates reflect the robustness of decoding performance of the members of the pools, while mean thresholds ideally reflect sensitivity to modulation. Because a threshold value is only assigned when a given pool exceeds a threshold criterion (see Methods) it is possible for recruitment effects to affect mean threshold values. In a recruitment effect, an improvement in success rate results in previously non-criterion pools reaching criterion – because these are likely to be pools with high modulation depth thresholds, their inclusion may result in mean threshold values that appear to be less sensitive, even if all individual pools improve (e.g. ref. 18, Figs. 6 and 7). For completeness, we reported both values, particularly because threshold estimates provide a more direct analogy to psychophysical performance metrics. Threshold measurements more commonly resulted in smaller effect sizes (e.g. Figs. 6 and 7) or noisy threshold estimates for decoding results based on phase locking (Fig. 10B, D), which could reflect the aforementioned recruitment effect. Overall, we believe that success rates likely provide a better reflection of the relative efficacy of the population models and have therefore emphasized success rates when discussing the implications of our results.

There are also important caveats to consider when comparing psychometric and neurometric thresholds. First, sufficient variation in decoding model performance, coupled with improvements in decoding for increasing population sizes, will typically provide a good match to psychophysical thresholds at a tested simulated population size (44). Assignment of the causally relevant pool size for the behavioral readout is not possible without causal manipulations that were not pursued in these experiments. Nevertheless, it is clear that population-based decoding models can reproduce the behavioral sensitivity observed in highly trained animal subjects with relatively small populations, and that the decoding models that best approximate behavioral thresholds are distinct in A1 and ML.

Our analyses relied on averaging the performance of randomly generated pools of each chosen size. Ince et al. (45) demonstrated that the population decoding performance at a given pool size can vary considerably depending on pool membership, which Downer et al. (20) confirmed for the decoding of sinusoidal AM signals similar to those employed here. Thus, the similarity between the neurometric and psychometric results must be understood in the context of the averaging process applied to the neurometric data, since the best and worst pools of a given size could span a significant range of thresholds. Relatedly, the pools were generated via random selection with replacement, so modulation frequencies that are less well represented in the dataset would exhibit less variation about the average, particularly for larger pool sizes. Therefore, the average result may not be representative of the true population average.

Even with sampling with replacement, the size of the pools did not completely asymptote, indicating that better decoding performance can likely be achieved with larger populations. However, the improvements expected for larger populations are potentially limited by correlations among the responding neurons (46–49). Downer et al. (20) demonstrated that population-based decoding of sinusoidal AM was worse for simultaneous recordings than for trial-shuffled versions of the same data, which remove noise correlations while retaining signal correlations. In the current study, all neurons included in a given pool were responding to an identical stimulus, but were recorded at different times. As such, the pools included signal correlations, but excluded ‘noise correlations’ that might reflect attention, decision-related activity, motor preparation, motor execution, or learning (50–53). Although noise correlations are believed to decrease encoding accuracy among neural populations with similar tuning (54–56), Johnson et al. (13) proposed that a two-pool opponent coding mechanism could make signal processing robust to changes in correlated noise among neurons (12), since increases in noise correlations have been shown to improve population-based encoding accuracy among neuronal pools with opposite tuning (46, 57–60). Thus, the emergence of bidirectional encoding of modulation depth in ML could be indicative of a processing strategy aimed at supporting dual encoding of sensory and cognitive-/task-related variables in higher auditory areas where the putative sources of noise correlations are expected to be more prominent. Understood in this context, “noise” correlations are not necessarily extrinsic to performance of the behavioral task, just the sensory demands of the task. For example, activity increases reflective of stimulus extrinsic factors that might be interpreted as increases in modulation depth among neurons with INC responses would be interpreted as decreases in modulation depth among neurons with DEC responses, thereby preserving stimulus feature information in the fuller cortical representation.

Unweighted, indiscriminate pooling obviates the need for ‘labeled line’ representations that track neuronal identity (the ‘labeling problem’; ref. 43). However, as noted above and elsewhere (22), heterogeneity in stimulus feature preferences corrupts rate-based encoding schemes for indiscriminate pools, just as heterogeneity in modulation phase preferences corrupts timing-based encoding schemes. Bidirectional encoding of modulation depth is especially problematic for indiscriminate rate codes, since it is analogous to populations of neurons responding in anti-phase in the context of timing-based codes. As such, the emergence and increasing prevalence of bidirectional AM encoding in the ascending auditory pathway is perhaps surprising. Unlike modulation transfer functions, which are heterogenous and often exhibit nonmonotonic tuning (1, 2, 61), modulation depth functions in subcortical and core cortical regions are typically monotonically increasing (2). Moreover, the significant reductions in synchrony to the modulation envelope among ML neurons (10, 11) constrains timing-based modulation encoding schemes (21, 22), which are more robust to stimulus context effects than firing rates (16).

What then is bidirectional encoding for, and, more broadly, what value do non-core auditory neurons add to modulation detection and discrimination, given their poorer synchrony and greater response heterogeneity? Our current work, understood in the context of prior work in the laboratory, is consistent with an important shift in how auditory information is represented across the auditory neuraxis. Lower processing centers, relative to higher processing centers, shield their representations from stimulus-extrinsic factors, likely because the flexibility of those representations is more tightly constrained by the need to optimize for efficient signal transmission. Higher centers, by contrast, are optimized for flexibility, in order to support arbitrary mappings to behavioral outputs, such as those required by experimental paradigms we have employed. We speculate that belt areas support integration of task-related, stimulus-extrinsic information, such that ‘noise correlations’, in the context of stimulus decoding, are better understood as multiplexed ‘signals’ related to task demands. As a result, emergent representations such as bidirectional encoding allow for complementary readout mechanisms that are more independent from ‘noise correlations’ in ML, relative to A1 (12). That said, bidirectional encoding is not an exclusive property of ML, but a manifestation contingent on specific task demands in highly trained animals, and can be found in A1 when task demands are different (38). The success of the two-pool opponent code in ML underscores the importance of readout mechanisms that can most effectively exploit cortical representations that balance stimulus-related and task-related information. Future work will be necessary to explore limits on the flexibility of rate-based representations of complex dynamic stimuli like AM, whether such flexibility necessarily comes at the price of the temporal fidelity observed in primary auditory cortex, and how such representations empower attentional mechanisms and deeper integration of stimulus-extrinsic, task-related activity.

## Author Contributions

MLS and KNO conceived and designed research; MN and KNO performed experiments; JSJ analyzed data and prepared figures; JSJ and BJM drafted manuscript; BJM, MLS, JSJ and KNO edited and revised manuscript.

## Acknowledgements

This work was funded by NIH NIDCD grant DC02514 (MLS)

